# Robustness of Collective Scenting in the Presence of Physical Obstacles

**DOI:** 10.1101/2021.03.23.436715

**Authors:** Dieu My T. Nguyen, Golnar Gharooni Fard, Michael Iuzzolino, Orit Peleg

## Abstract

Honey bees (*Apis mellifera* L.) aggregate around the queen by collectively organizing a communication network to propagate volatile pheromone signals. Our previous study shows that individual bees “scent” to emit pheromones and fan their wings to direct the signal flow, creating an efficient search and aggregation process. In this work, we introduce environmental stressors in the form of physical obstacles that partially block pheromone signals and prevent a wide open path to the queen. We employ machine learning methods to extract data from the experimental recordings, and show that in the presence of an obstacle that blocks most of the path to the queen, the bees need more time but can still effectively employ the collective scenting strategy to overcome the obstacle and aggregate around the queen. Further, we increase the complexity of the environment by presenting the bees with a maze to navigate to the queen. The bees require more time and exploration to form a more populated communication network. Overall, we show that given volatile pheromone signals and only local communication, the bees can collectively solve the swarming process in a complex unstructured environment with physical obstacles.

## 1 Introduction

Animals in large groups must effectively communicate to exchange information and coordinate group processes. Volatile chemical signals (i.e. pheromones) are a prevalent communication signal in the honey bees, crucial to the colony’s coordinated processes, such as foraging and caste recognition [1, 6, 11]. Individual bees use their antennae to receive and respond to specific chemical signals, and transmit different pheromones in particular contexts. A limitation of the pheromone signal lies in its spatiotemporal decay, which limits the range and timing of information exchange. In the scenario of worker bees aggregating into a coherent swarm around the queen by communicating via pheromones, we previously showed that the bees form a signal propagation strategy to create a communication network that overcomes the limitation of the decaying signals [8]. In this communication network, individual bees sense local pheromone gradients above a concentration threshold and “scent” [7, 9]: They raise their abdomens to release pheromones from the Nasonov gland and fan their wings to direct the pheromone diffusion behind them and disperse the signals to other bees. This directional bias allows bees far away from the queen to sense the amplified signals and further propagate them. These pheromone detection and transmission events create a dynamic communication network that allows worker bees to localize the queen and effectively aggregate around her into a coherent swarm.

In this work, we extend our previous study on the mechanisms of the honey bee aggregation [8] by introducing environmental stressors. Honey bees form swarms in variable and unpredictable landscapes. While studies have shown the ability of insects to navigate through unfamiliar environments to reach food rewards [12, 13], we explore how honey bees conduct collective olfactory communication to localize the queen and swarm. We present the bees with physical obstacles (i.e., long bars) that partially block pheromone signals and prevent a wide path to the queen. Further, to test the effect of a more challenging obstacle, we present the bees with a maze. Through these experiments, we aim to assess whether the honey bees’ collective scenting communication can overcome stressors that increase the complexity of the swarming process, and how the difficulty level of those stressors affect the bees’ ability to collectively localize the queen.

## 2 Methods

### 2.1 Experimental setup

We restrict our backlit arena (50 cm × 50 cm × 1.5 cm) to be semi-two-dimensional to prevent flying, as bees have been shown to scent while standing [7]. We record the bees from an aerial view with a video camera (4k resolution, 30 fps). Before the experiments, worker bees are isolated from their queen for 24 hours and then introduced to a caged queen (10.5 × 2.2 × 2.2cm) from our queen bank for another 24 hours. To begin an experiment, the caged queen is isolated from the workers and placed into the arena. We use wooden obstacles that are the height of the arena to ensure bees cannot climb over them. Workers are then placed into the arena, and the plexiglass is placed on top to enclose the space. Temperature is monitored regularly to ensure that the heat from the backlight board stays below 32 − 35°C and does not affect the fanning behavior.

### 2.2 Bee detection & scenting recognition

We employ computer vision and deep learning approaches to automatically detect scenting bees and estimate their orientation, as originally presented in detail in [8]. To detect individual bees in the videos, we extract images at 30 fps and use Otsu’s method to adaptively threshold the images [10]. We iteratively apply morphological transformations (opening) to remove noise and separate individual bees from clustered groups [2]. Connected components is then applied to obtain the components’ centroids (x,y positions) and areas. Using the component areas, we filter out large clusters to isolate the individual bees.

To classify individual bees as scenting or non-scenting, we train a ResNet-18 convolutional neural network (CNN) model [5] using 28,458 labeled images [8]. The model is trained with data augmentation (horizontal and vertical flipping, brightness adjustments, scaling, translation, and rotation) and balanced sampling to combat the class imbalance (9:1 non-scenting to scenting) for 1203 epochs with early stopping to prevent overfitting. On the test set, we achieve 95.17% accuracy, indicating that our model can generalize to unseen data.

We use the same ResNet-18 architecture for orientation prediction. The loss function is modified for the model to output continuous values for the predicted angles: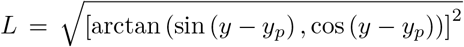, where *y* is the true label, and *y*_*p*_ is the network’s prediction. We created a labeled dataset of 15,435 images, each with head and tail positions, from which we compute the ground-truth orientation angle. On the test set, this model achieves 96.71% with 15° of error tolerance.

### 2.3 Analysis of time-series data

With the positions and orientations of scenting bees, we obtain several time-series properties from the experiments. The number of scenting bees are extracted per frame, presented as a rolling mean with the window size of 100 frames. We also extract the average distance to the queen. Because our bee detection method cannot detect every single individual bee when they touch or overlap, we compute the average distance of all black pixels to the queen’s location. This is an effective method to measure the distance to the queen over time, as the queen’s cage and the obstacle are stationary, and the remaining black pixels in the arena only represent the moving worker bees. For each of these properties, we average the time-series data across all five density experiments for every condition (no obstacles, bar, maze) and obtain the standard deviation.

### 2.4 Attractive surface reconstruction

We then correlate the scenting events with the spatiotemporal density of bees. For each scenting bee *i* at time *t*, we define its position as 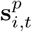, and its direction of scenting as 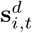 (unit vector). Assuming the scenting bees provide directional information to non-scenting bees, we treat 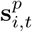 and 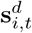 as a set of gradients that define a minimal surface of height *f* (*x, y, t*). Thus, *f* (*x, y, t*) corresponds to the probability that a randomly moving non-scenting bee will end up at position (*x, y*) by following the scenting directions of scenting bees:

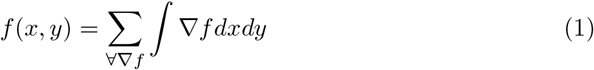

where 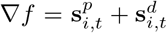. We regularize the least squares solution of the surface reconstruction from its gradient field, using Tikhonov regularization [3, 4].

To obtain the time at which the attractive surface first correlates to the final clustering of the bees (i.e., when the area around the queen becomes maximal in the surface), we treat the surface as an image to segment out the region that is above a particular threshold, computed from obtaining the image histogram and its peak. The surface becomes a binarized image of foreground and background pixels. The connected component algorithm is applied to the binarized image to isolate the regions of high values in the surface. We define a square region (850×850 pixels in a 1800×1800 pixel image) around the queen and check for the proportion of that region being covered in the resulting surface (e.g. 50%).

## 3 Results

### 3.1 Linear Obstacles

To characterize how a physical obstacle affects the aggregation process of the bees, we compare the dynamics when an obstacle is present and when it is absent (control). We present the obstacle as a long wooden bar that lies diagonally in the middle of the arena, blocking most of the pathway from the worker bees to the queen in the cage. In both conditions (control and the bar), the caged queen bee is placed at the top-right corner of the arena, while workers bees are placed at the bottom-left. For each condition, we performed five experiments with densities ranging from 205 to 420). In Fig. 2A, we show snapshots and the moving average attractive surfaces for an example control experiment (*N* = 320). Over approximately 1800 seconds (~ 30 minutes), the bees search for the queen and aggregate around her. The bees activate a scenting network early on in the experiment, as shown in the snapshot at *t* = 140 sec. Here, the surface begins to reflect how the collective directional scenting events point the bees to aggregate around the queen’s area (i.e. regions of higher values). Most of the swarm around the queen forms at *t* = 900 sec (~ 15 min). In Fig. 2B, we show snapshots and the moving average attractive surfaces for an example linear obstacle experiment (*N* = 310). Here, the bees search around the space behind the bar until a few bees find the opening and scent to inform other bees as they collectively make their way to the queen, at around 900 sec (~ 15 min). Most of the worker bees swarm around the queen by *t* = 1800 sec (30 min).

**Fig. 1:**
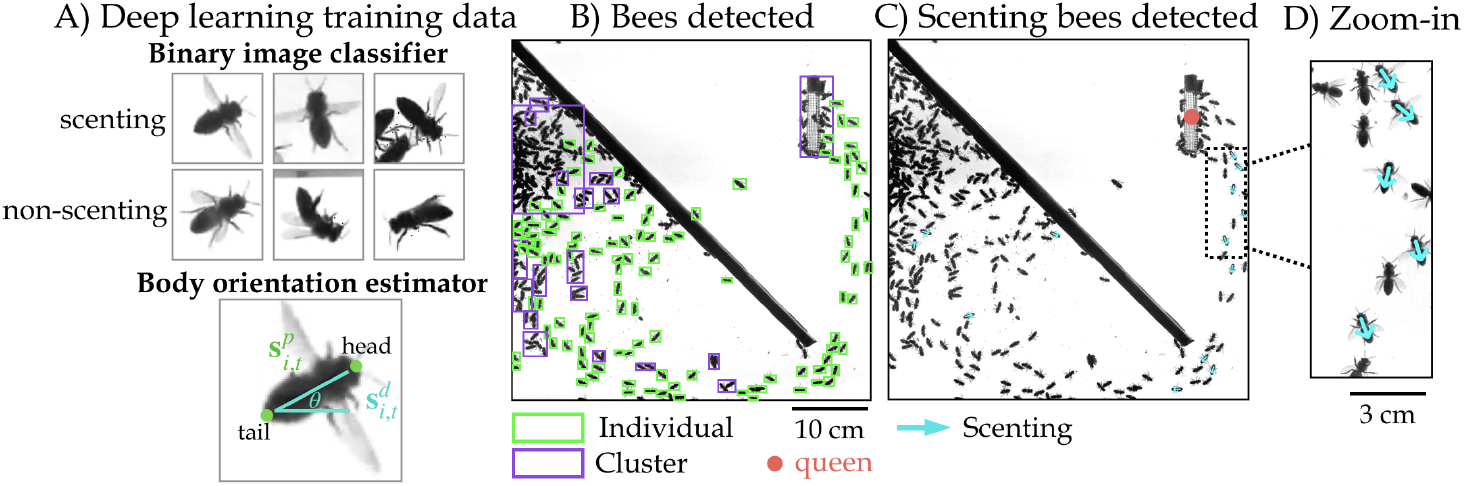
Bee detection and scenting recognition. A) Training data examples for two deep learning models, the binary image classifier to identify scenting and non-scenting bees and the orientation estimator. B) Example detections of individual bees (green) and clusters (purple). The largest cluster containing the obstacle is not shown to reduce visual clutter. C) Example detections of scenting bees and their orientations (teal arrows). D) A zoom-in showing example scenting bees with wide wings.

**Fig. 2:**
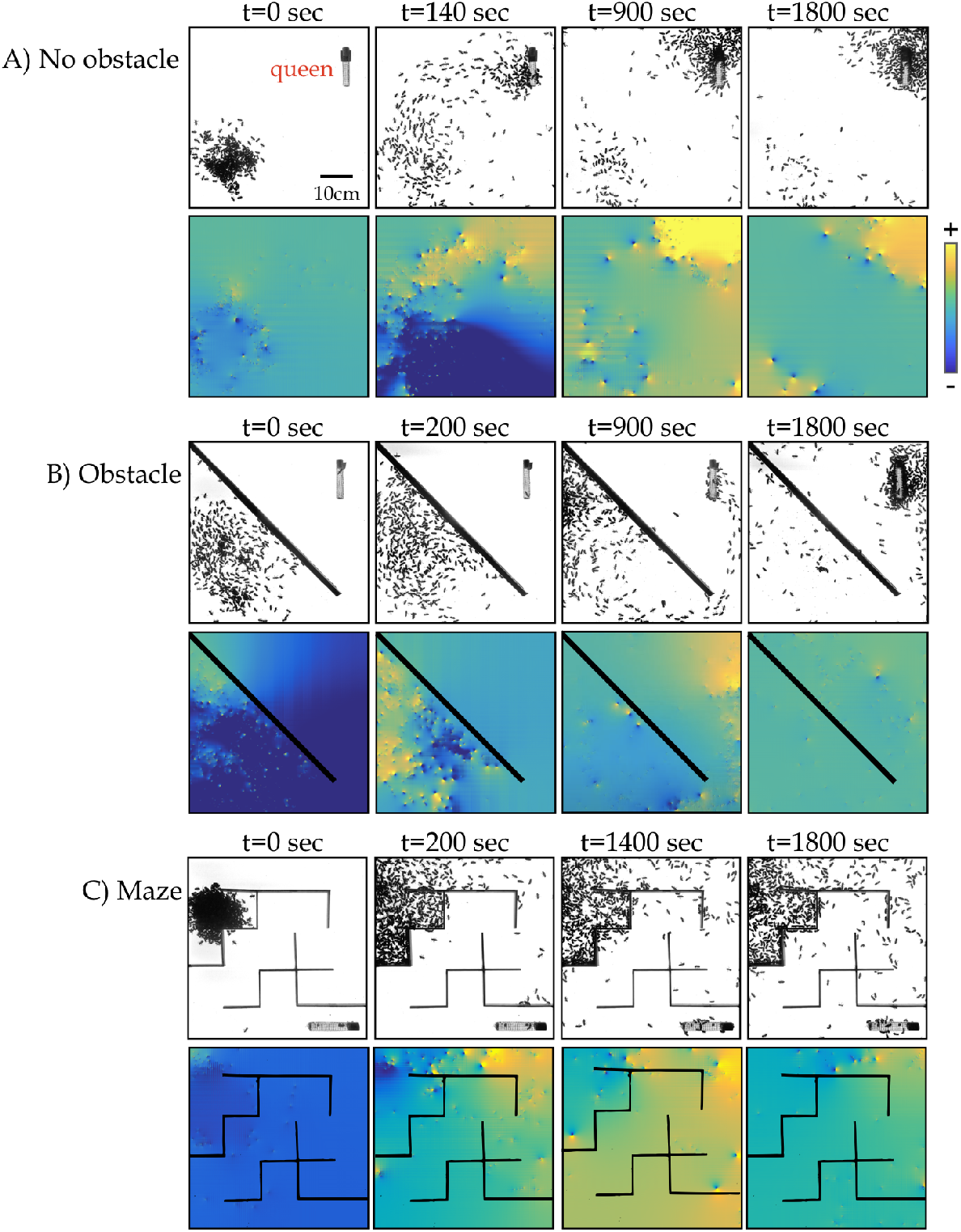
Snapshots and attractive surfaces of three experimental conditions. A) Snapshots of a control experiment where the bees are in a semi-2D area with a caged queen, without any physical obstacles. Over time, the bees form the communication network with collective scenting and cluster around the queen. The corresponding attractive surfaces *f* according to Eq. 1 are shown below each snapshot to show how the collective scenting events correlate to the spatial-temporal density of bees. B) An experiment where the bees are placed on one side of a bar obstacle. C) An experiment where the bees are placed in a maze with a caged queen at the end of the correct path.

### 3.2 Maze

To determine whether a more complex physical obstacle affects aggregation, we also present the bees with a maze to navigate and find the queen. In Fig. 2C, we show snapshots and the moving average attractive surfaces for an example maze experiment (*N* = 450). Similar to in control and linear obstacle experiments, we place the caged queen in a far corner from the worker bees. Here, some workers explore the paths in the maze and scent as they escape from the initial corner, at around *t* = 200 sec. At *t* = 1400 sec, more bees follow the scent to escape and some are able to find the queen. Compared to the control condition (Fig. 2A) and a simple bar obstacle (Fig. 2B), the maze is noticeably more difficult for most of the bees to escape the initial corner and aggregate around the queen.

### 3.3 Quantitative comparisons of the different conditions

We quantitatively compare the aggregation process in the three different conditions by observing the number of scenting bees over time (averaged over the five experiments per condition, with standard deviation shown as the shaded area), as shown in Fig. 3A. Without an obstacle, the sharp peak in the number of scenting bees occur early on in the experiments (green curve) and gradually decreases as most bees cluster around the queen. With a bar obstacle (red curve), the peak also occurs very early on, but behind the bar as the bees have yet to find the escape opening. A small second peak occurs at approximately ~900 sec, when the bees find the opening and begin to cluster around the queen. In the maze experiments (orange curve), the number of scenting bees is relatively stable and low over time. Overall, we observe significantly more scenting bees in the absence of obstacles, and less scenting bees with increasingly difficult obstacle.

**Fig. 3:**
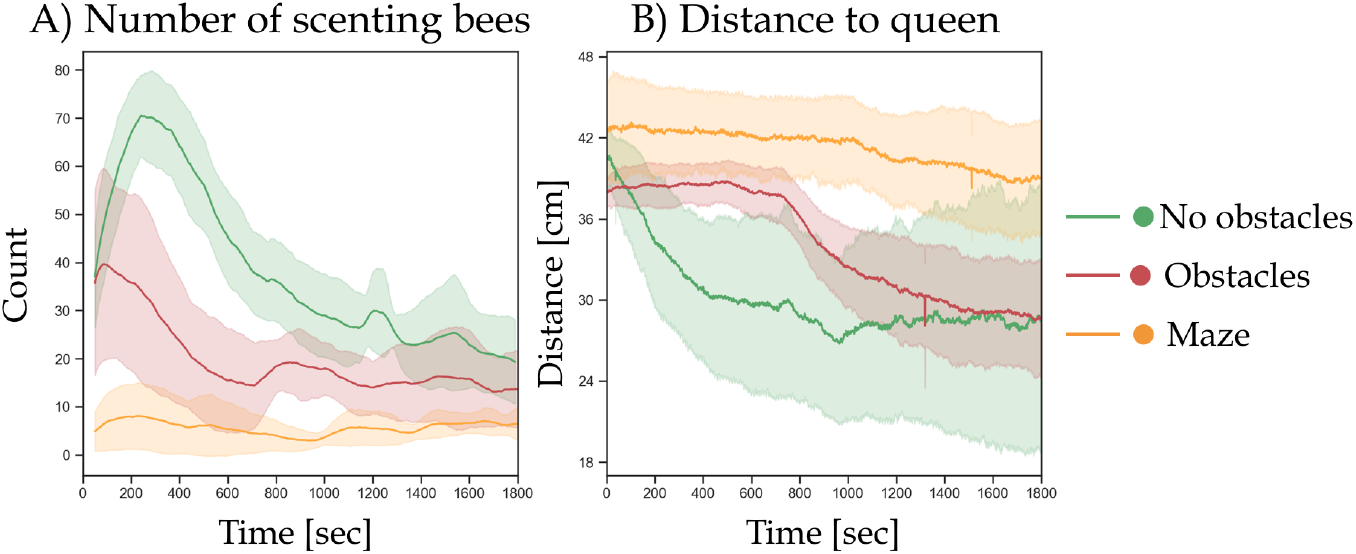
Time-series data of bee aggregation in three conditions, with no obstacles (green), a linear obstacle (red), and a maze (orange). A) The average number of scenting bees over time across five trials per experimental condition. B) The average distance to the queen over time across five trials per experimental condition.

We also extract the average distance to the queen over time Fig. (3B). As shown, the distance sharply decreases early on in the control experiments (green curve), as the bees quickly activate the scenting network and begin clustering around the queen. On the other hand, with the bar obstacle (red curve), the bees spend some time at the beginning exploring the complicated space and finding the escape opening, reflected in a relatively flat curve until ~850 sec, when they find the opening and therefore a path to the queen as the average distance to the queen decreases. In the maze experiments (orange curve), given the more complex obstacle, we observe a relatively flat average distance to the queen until a slight decrease at ~1100 sec, when a small fraction of the bees escape the initial corner of the maze and find the queen.

Finally, we extract the time at which the attractive surface begins to correlate to the clustering of the bees in the queen’s area. We define 50% coverage of the square region around the queen’s position as the threshold at which we extract this time point. These time points for all experiments across three conditions are presented in Table 1. Compared to the control condition, the time to reach the threshold coverage of the queen’s area is significantly higher when the bees must navigate a linear obstacle or a maze.

**Table 1:** Time to reach 50% coverage (TC) for all experiments.

| Experiments | No Obstacles |  |  |  |  | Obstacles |  |  |  |  | Maze |  |  |  |  |
| --- | --- | --- | --- | --- | --- | --- | --- | --- | --- | --- | --- | --- | --- | --- | --- |
| Density | 250 | 310 | 320 | 370 | 400 | 205 | 240 | 310 | 380 | 400 | 100 | 115 | 415 | 460 | 450 |
| (TC)(sec.) | 320 | 20 | 80 | 60 | 180 | 820 | 1060 | 820 | 880 | 960 | 1820 | 1380 | 2800 | 1880 | 1400 |
| Average TC (sec.) | 132 |  |  |  |  | 908 |  |  |  |  | 1856 |  |  |  |  |
| Std Dev TC (sec.) | 107.77 |  |  |  |  | 91.74 |  |  |  |  | 515.27 |  |  |  |  |

## 4 Conclusion & Discussion

Extending our previous study on the communication mechanisms that honey bee swarms use to locate their queen [8], we introduce environmental stressors in the form of physical obstacles to the system. Worker bees are placed into an arena with physical obstacles that partially block pheromone flow and prevent a wide, open path to the queen. We find that similar to the control experiments without any obstacles, the bees still employ the scenting behavior to locate the queen and propagate the signals, as shown in our attractive surface analysis. However, given the physical obstacles, the bees require more time to find the escape points to break free from the initial position and swarm around the queen. Moreover, as the difficulty of the obstacle increases, from a bar to a maze, there are significantly less scenting events that may result in the smaller fraction of the worker bees being able to find the queen. Overall, we show that although pheromone signals are volatile and communication is local on the individual level, the bees require more exploration and time to navigate their way around the obstacles, but can ultimately collectively handle a more complex environment.

For future works, to better understand the communication mechanisms specific to the swarming scenario with physical obstacles, we will also introduce obstacles to our agent-based model of the honey bee swarming phenomenon presented in [8]. The model will allow us to explore how certain behavioral parameters, such as the pheromone detection threshold and the magnitude of the wing fanning to disperse pheromones directionally, vary as the environment changes and becomes more complex. Furthermore, as physical obstacles are not the only environmental stressors honey bees may encounter during the swarming process, we will also explore other stressors to test the adaptability and resilience of the pheromone communication network. For example, we will introduce artificial pheromones that act as a secondary signal that interferes with the queen’s signals or disrupt pheromone flow with wind at various wind speeds. Our current experimental and modeling tools will allow us to conduct these extensions and gain a deeper understanding of how the honey bees find creative solutions to effectively communicate and achieve their collective goals. The understanding of the behavior of this biological system and their adaptive solutions can inspire designs and improvements in non-biological systems in which individuals are limited to local interactions but contribute to a coordinated collective process.

## Acknowledgments

This work was supported by the National Science Foundation Graduate Research Fellowship under Grant No. DGE 1650115 (D.M.T.N.) and Physics of Living Systems Grant No. 2014212 (O.P.). Any opinion, findings, and conclusions or recommendations expressed in this material are those of the authors(s) and do not reflect the views of the NSF. We also acknowledge the BioFrontiers Institute (internal funds), the Interdisciplinary Research Theme on Autonomous Systems (O.P.).

